# First insights into nasal microbiome in wine tasters

**DOI:** 10.1101/2023.03.21.533426

**Authors:** Sofia Duarte-Coimbra, Giovanni Forcina, Lucía Pérez-Pardal, Albano Beja-Pereira

## Abstract

Over the last decades, the study of the microbiome has been receiving increasing attention as a major driver of individual health and wellbeing. The accumulation of knowledge on microbiomes sparked new research lines, from which the association between oral microbiome composition and taste perception is of great interest. Taste plays a paramount role in food and beverage choice as well as emotions. For wine tasters, the smell is also part of the tasting evaluation. However, the nasal microbiome is relatively unexplored. The relation between the microorganisms residing in the nostrils is still poorly known despite their leading role in flavor perception. Therefore, characterizing the composition of nasal microbiomes represents a fundamental prerequisite to elucidate their relationship with taste. To improve our understanding of the relationship between taste and the microorganism inhabiting the nostrils, the nasal microbiome of 5 wine tasters versus 5 non-tasters was analyzed through the sequencing of the V3-V4 region of the *16S rRNA* gene. The taxonomic composition of these nasal microbiomes was characterized, and the comparison of diversity indexes revealed no significant differences. However, the experimental group showed a lower number of identified taxa (171) when compared to the control group (287). Another interesting result was the higher presence of Krebs Cycle pathways in wine tasters, which could indicate the importance of the nostril bacterial community in alcohol oxidation. Regarding smoking habits, smokers presented a lower microbiome diversity. These preliminary results should be confirmed in a larger sample dataset of wine tasters and controls.

## Introduction

Wine tasting is a multimodal perception arising from the functional integration of data obtained through the chemical senses, namely gustation and olfaction, with oral and nasal somatosensory inputs. Flavor sensation changes among individuals, with part of this variation being likely attributable to differences in oral microbiota composition (Schwartz et al., 2021).

Although relatively unexplored, the microbiome of the nasal cavity is deemed to play a leading underlying the connection between taste and smell (Small & Green, 2012). Firmicutes and Actinobacteria - the two most represented bacterial phyla in the oral cavity - are also those dominating nostril microbiomes (Bassis et al., 2014). Even though the relationship between taste perception and nasal microbiota composition has not been investigated in depth, it was hypothesized that microorganisms could modulate the olfactory function by generating strong odor products (Koskinen et al., 2018).

In resume, the influence of nasal microbiome composition on wine tasting might reveal how continuous exposure to wine modulates the sensorial capacities of an individual. The present study aimed to elucidate the relationship between taste perception and nasal microbiome composition, as well as possible individual differences in the community of microorganisms residing in the nostrils. To achieve these goals, we amplified and sequenced the hypervariable region V3-V4 of the 16s *rRNA* gene in the nasal microbiome of 5 wine tasters and 5 individuals from a control group. The influence of other factors such as sex and lifestyle were also evaluated concerning microbiome composition.

## Materials and methods

### 1. Sampling and questionnaire administration

A total of 5 volunteers were recruited from Bairrada Viticulture Commission and oenologists from Coimbra (central Portugal) to form the experimental group of wine tasters (age range 43 to 63; average 51). The control group was constituted of 5 volunteers (age range 40 to 53; average 48) not related to wine professional activities (Table 1). The oral microbiome of these 10 individuals was characterized in a previous study (Duarte-Coimbra et al., 2023), during which nasal samples were collected simultaneously. The participants were asked not to eat, drink, smoke, brush their teeth and chew gums for two hours prior to sample collection. Volunteers that had been under the effect of antibiotics over the last month prior to the survey were refused. The participants employed swabs (SK-2S Isohelix Swab, Harrietsham, UK) to collect nasal samples by introducing a sterile swab inside the nostril and rotating it for about 60 seconds against the nasal wall. Samples were preserved in RNAlater (Thermo Fisher Scientific, Waltham, MA, USA) and stored at −20°C until DNA extraction. All samples were collected after reading and signing up the informed consent. Sample collection was performed by the volunteers themselves who later filled out an online questionnaire via Google Forms (Google, Mountain View, California). Personal information about food habits, lifestyle, and oral hygiene was collected. Additionally, aspects of the professional activity of wine tasters were also requested (Supplementary Table 1). This research was authorized by the Bioethics Committee of CIBIO/InBIO (University of Porto).

**Table 1.**
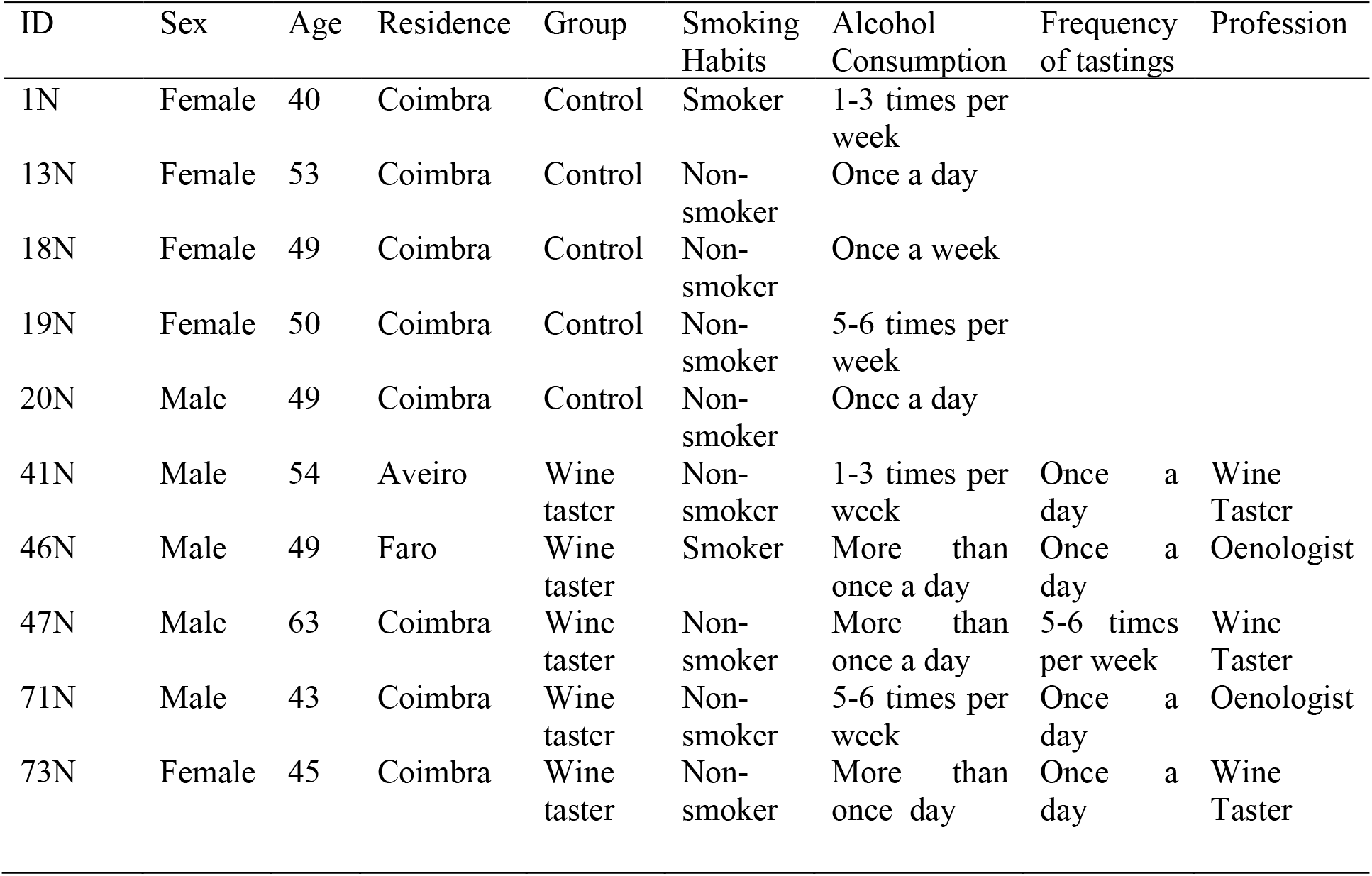
Characterization of study participants.

### 1. DNA extraction

DNA was isolated with the MagMAX Microbiome Ultra Nucleic Acid Isolation Kit (Thermo Fisher Scientific) following manufacturer’s instructions with minor modifications, particularly by adding 1μl RNase (4ng/μl) at each one of the two washing steps and eluting the DNA in 50μl Elution Buffer. One extraction blank was incorporated to detect possible contamination. DNA was quantified with a Qubit™ dsDNA HS Assay Kit (Thermo Fisher Scientific).

### 2. *16S rRNA* gene amplification, library preparation and sequencing

A two-step Polymerase Chain Reaction (PCR) approach was used to amplify the V3-V4 hypervariable regions and index individual samples prior to pooling and sequencing following Almeida-Santos et al. (2021). PCR products were run on a 2% agarose gel to check for the expected approximate size of 650 bp. PCR blanks were incorporated to check against possible contamination. Amplicons were cleaned up with 0.8X AMPure XP beads (Beckman Coulter, Brea, CA, USA) and two 80% ethanol washes prior to resuspension in 25 μl EB buffer (Qiagen, Venlo, the Netherlands). Finally, individual libraries were quantified using a BioteKTM EpochTM Microplate Spectrophotometer (Thermo Fisher Scientific). All samples were normalized to 20 nM and pooled.

Pool quantification and sizing were implemented in a TapeStation 2200 (Agilent Technologies, Santa Clara, CA, USA). The pooled library was exported to NovoGene (Cambridge, UK) and sequenced with a 2×250bp paired-end configuration on an Illumina MiSeq platform applying the MiSeq v2 500-cycle sequencing kit (Illumina Inc., San Diego, CA, USA).

### 3. Sequence processing and alignment

The raw paired-end sequences data were analyzed with QIIME (Quantitative Insights into Microbial Ecology) v.2-2021.4 (Bolyen et al., 2019) and processed with quality control package DADA2 (Callahan et al., 2016) using the successive steps: filtering, denoising, dereplication, chimera identification, and merging. SILVA database version 138 (Quast et al., 2013) was used to blast against the obtained Amplicon Sequence Variants (ASV), thus obtaining a table of ASV frequency per sample based on the Naïve Bayes classifier trained on the SILVA database.

QIIME2 was also used to perform taxon identification. Mitochondrial and chloroplast ASVs were excluded from downstream analyses.

### 4. Statistical analyses

ASV relative frequencies and taxonomic abundance were calculated for each sample with QIIME2 for phylum, class, genera, and species level. The differences between the experimental and control group were calculated using the Mann-Whitney U test in SPSS v.27.

Alpha diversity, incorporating the richness (number of distinguishable taxa) and evenness (taxa distribution) within a given sample, was estimated employing two metrics: ASV abundance and Shannon diversity index (SDI) (Shannon, 1948). Beta diversity, indicating differences in the taxonomic structure between samples, was estimated through Bray-Curtis dissimilarity (Bray & Curtis, 1957). Rarefaction curves were performed to assess sampling depth for the diversity analyses.

Alpha and beta diversity were compared using a nonparametric Kruskal-Wallis H test and a Permutational Multivariate Analysis of Variance (PERMANOVA) (Anderson, 2001) with 999 permutations, respectively, in QIIME2. To investigate the putative clustering of the samples analyzed, we carried out a Principal Coordinated Analysis (PCoA) in EMPeror (Vázquez-Baeza et al., 2013) using the Bray-Curtis matrix. Finally, heatmaps were generated and hierarchical clustering analysis performed using the R package *phyloseq* (Fukuyama, 2020).

The variables analyzed by the dispensed questionnaire were also compared with the data obtained in QIIME2 analysis. Regarding the dietary habits of both wine tasters and non-tasters (n=10), different foods were classified according to their flavor (astringent, bitter, sweet, and sour) and consumption frequency. To compare the variables, we used the mean frequency per individual and five categories (Category 1: 0-20%; Category 2: 20-40%; Category 3: 40-60%; Category 4: 60-80%; Category 5: 80-100%) for a better understanding of data distribution (Supplementary table 1). We tested the occurrence of significant differences between the two groups based on a Crosstab with a Chi-Square T-test in SPSS. The same test was also applied to evaluate condiment consumption frequency.

Concerning oral hygiene, smoking habits, and alcohol consumption, we applied a descriptive analysis based on a Crosstab with a Chi-Square T-test in SPSS to explore the occurrence of significant differences between groups with regard to microbiome composition as well as possible associations between the variables and either the experimental or the control group. We also investigated the occurrence of specific associations between the experimental group and career length, frequency of tastings, number of tastings in the last 15 days and the most frequently tasted wine by means of Spearman correlation in SPSS. Finally, a Spearman correlation was also performed to investigate putative associations between lifestyle, dietary, and oral hygiene habits with the taxa obtained.

### 5. In-silico Prediction of Functional Composition

We employed PICRUSt2 *in silico* method (Phylogenetic Investigation of Communities by Reconstruction of Unobserved States) (Douglas et al., 2020) to predict the metagenomic functional composition of the nasal microbiome and STAMP (Statistical analysis of taxonomic and functional profiles) v.2.1.3 graphical software package (Parks et al., 2014) to compare the predicted pathways between wine taster and control groups based on a Welch’s two-tailed *t*-test.

## Results

### 1. Sequencing

A total of 1,705,517 reads were obtained with a mean of 155,047 reads per sample (range: 99,695-289,561). After quality filtering, 1,423,397 reads were retained. Extraction and PCR blanks produced no reads.

The experimental group yielded a lower average number of reads per individual (99,695) than the control group (115,228). High-quality reads were assigned to 1,188 ASVs with a total absolute frequency of 275,051 reads. The median ASV frequency per sample was 24,890 (range: 13,417–35,221). After sample rarefaction (threshold of 20,270), 202,700 reads (73.70%) and 10 samples were retained.

### 2. Microbiome characterization and taxonomy assignment

ASVs blast against the SILVA database turned into the identification of 361 taxa at the genus (87%) or species (42%) level, with a small fraction (5%) of the reads remaining unidentified (Supplementary table 2). The experimental and control groups shared 97 taxa, with the latter hosting 190 taxa not found in the experimental group, which, in turn, presented 74 private taxa. Overall, the nasal microbiome was dominated by phyla Actinobacteria (57%), Firmicutes (35%), Proteobacteria (8%), and Bacteroidota (1%). Among the 15 taxa accounting for the highest number of reads, the genus *Corynebacterium* was the most abundant.

After sequencing, processing, and blasting, we detected a total of 23 phyla, 32 classes, 79 orders, 135 families, 255 genera, and 152 species.

The mean taxa distribution of each individual was 4 phyla, 6 classes, 12 orders, 18 families, 55 genera and 20 species. *Corynebacterium* was the most frequent genus of the 10 most abundant genera (Figure 1).

**Figure 1.**
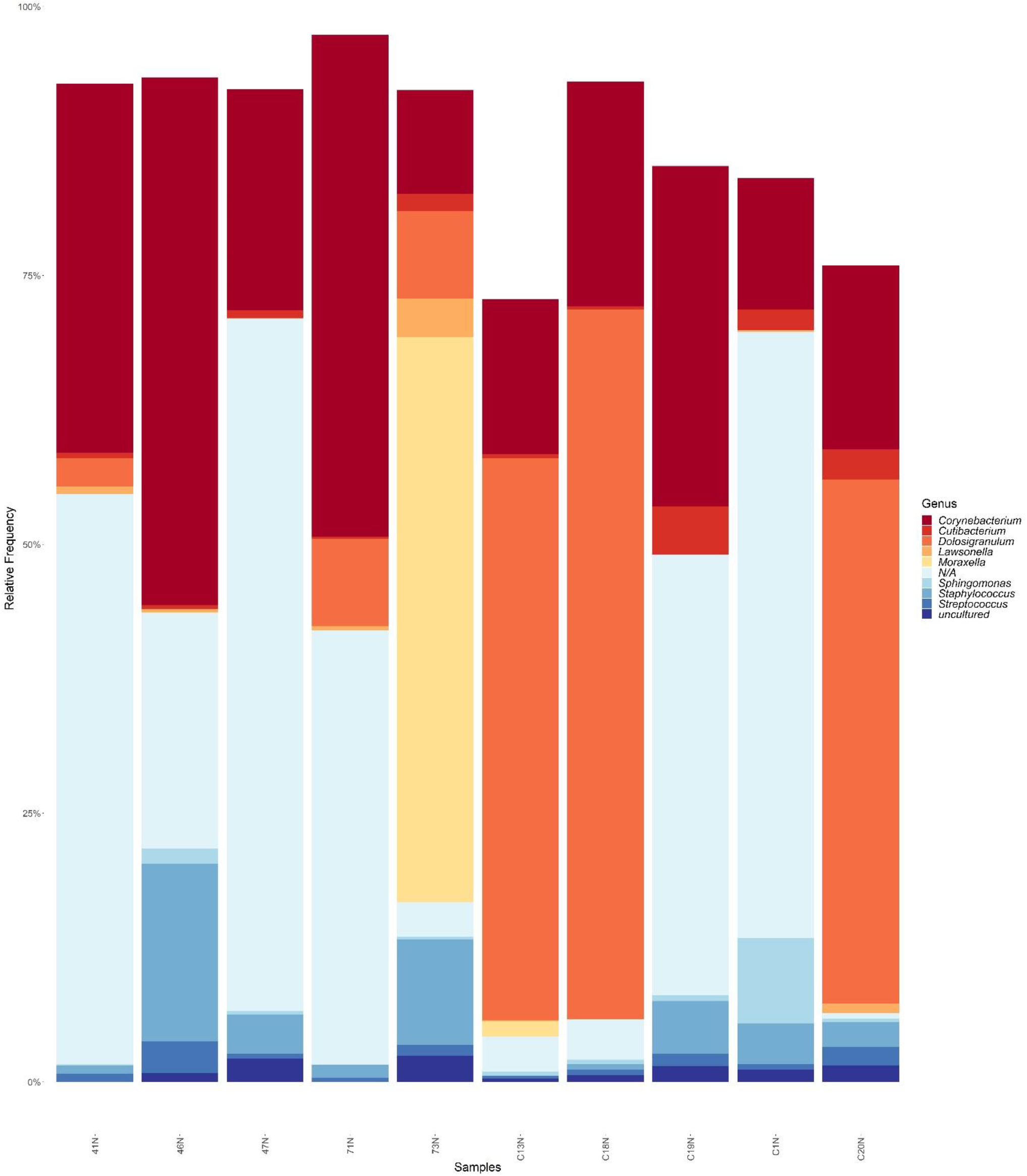
Relative abundance of the top 10 bacterial genera in nasal microbiome of samples belonging to the experimental and control group per individual. Sample names starting with C are from the control group.

Taxa distribution at the class level did not show significant differences between the experimental and control groups. However, while the relative frequency of Actinobacteria was higher in the wine taster group, Bacilli were higher in the control group. The dominant classes (accounting for 97%) that were present in more than 15% of the individuals belonging to either group were Bacilli, Alphaproteobacteria, Actinobacteria and Gammaproteobacteria (Figure 2A).

**Figure 2.**
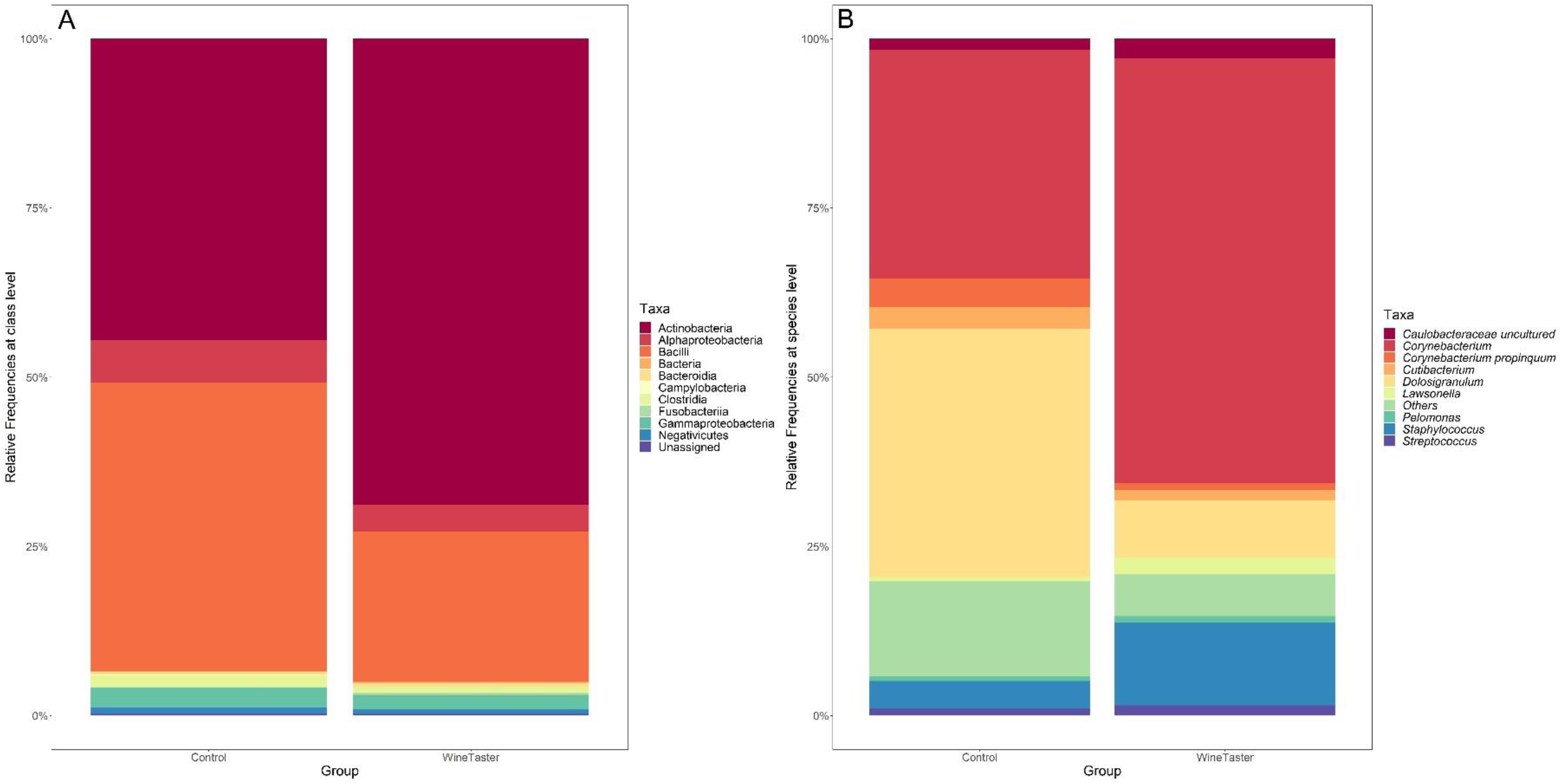
Taxa up to the class (A) and species (B) level identified in nasal microbiome samples of the experimental and control group.

No significant differences were found between groups when comparing the relative frequencies of the species identified. The ten most abundant species represented approximately 85% to 93% of the total in the experimental and the control group, respectively, with *Corynebacterium* being the most abundant genus in both groups. However, while this taxon was more copious in the experimental group, the control group was dominated by *Dolosigranulum* (Figure 2B).

Two heatmaps were constructed based on the relative abundances, one based on all the 361 taxa up to the genus level, and another with the taxa present in more than 15% of the samples (Figure 3). *Corynebacterium* and *Dolosigranulum* were the most abundant taxa, originating two major clusters based on their relative abundances, each one dominated by either one or the other genus. Interestingly, the cluster dominated by *Dolosigranulum* was entirely composed of individuals from the control group.

**Figure 3.**
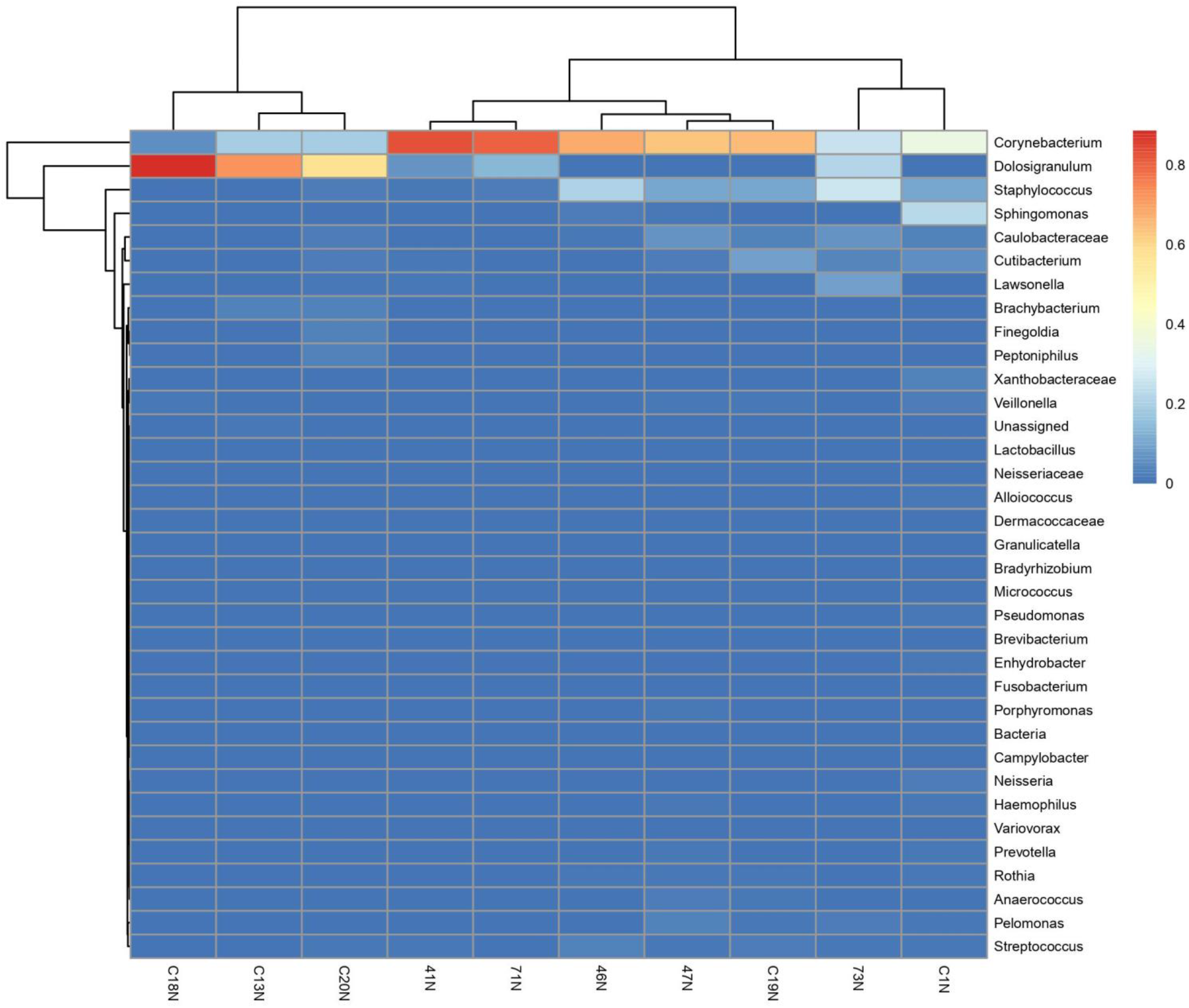
Hierarchical cluster of the most abundant genus per subject. Sample names starting with C are from the control group.

### 3. Diversity measures

Alpha diversity: The SDI estimates were 5.22 ± 0.22 and 5.67 ± 0.41 for the experimental and control group, respectively, whereas ASV abundance was 130.2 ± 21.99 and 200.4 ± 56.51 following the same order. No significant differences in the distribution of the SDI nor of ASV abundance were observed between groups (Figure 4A).

**Figure 4.**
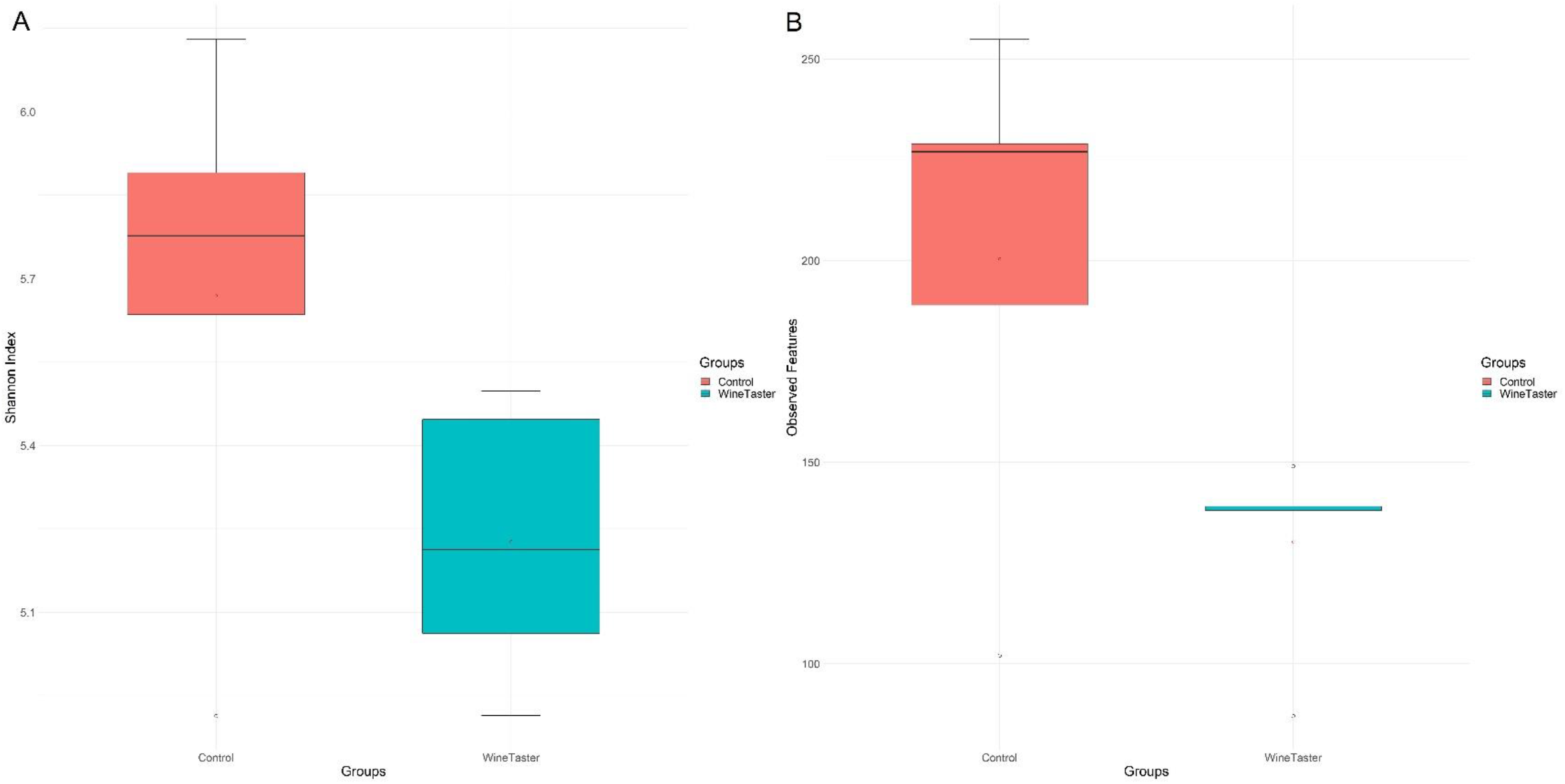
Boxplot charts depicting the distribution of Shannon Diversity Index (A) and ASV abundance (B) in the experimental and control group. Red dots represent the mean of each group.

Beta diversity: The distribution of pairwise Bray-Curtis dissimilarity values across individuals within and between groups evidenced no dissimilarities (Figure 4B). This pattern was confirmed by the PERMANOVA analysis (*pseudo-F*=1.867; *p*=0.055) and the unweighted UniFrac distances (*pseudo-F*=1.237; *p*=0.134), indicating no significant differences in the nasal microbiome composition of the experimental and the control group.

The PCoA revealed four clusters, two of which consisted of individuals belonging to the experimental group and the other two from the control group (Figure 5).

**Figure 5.**
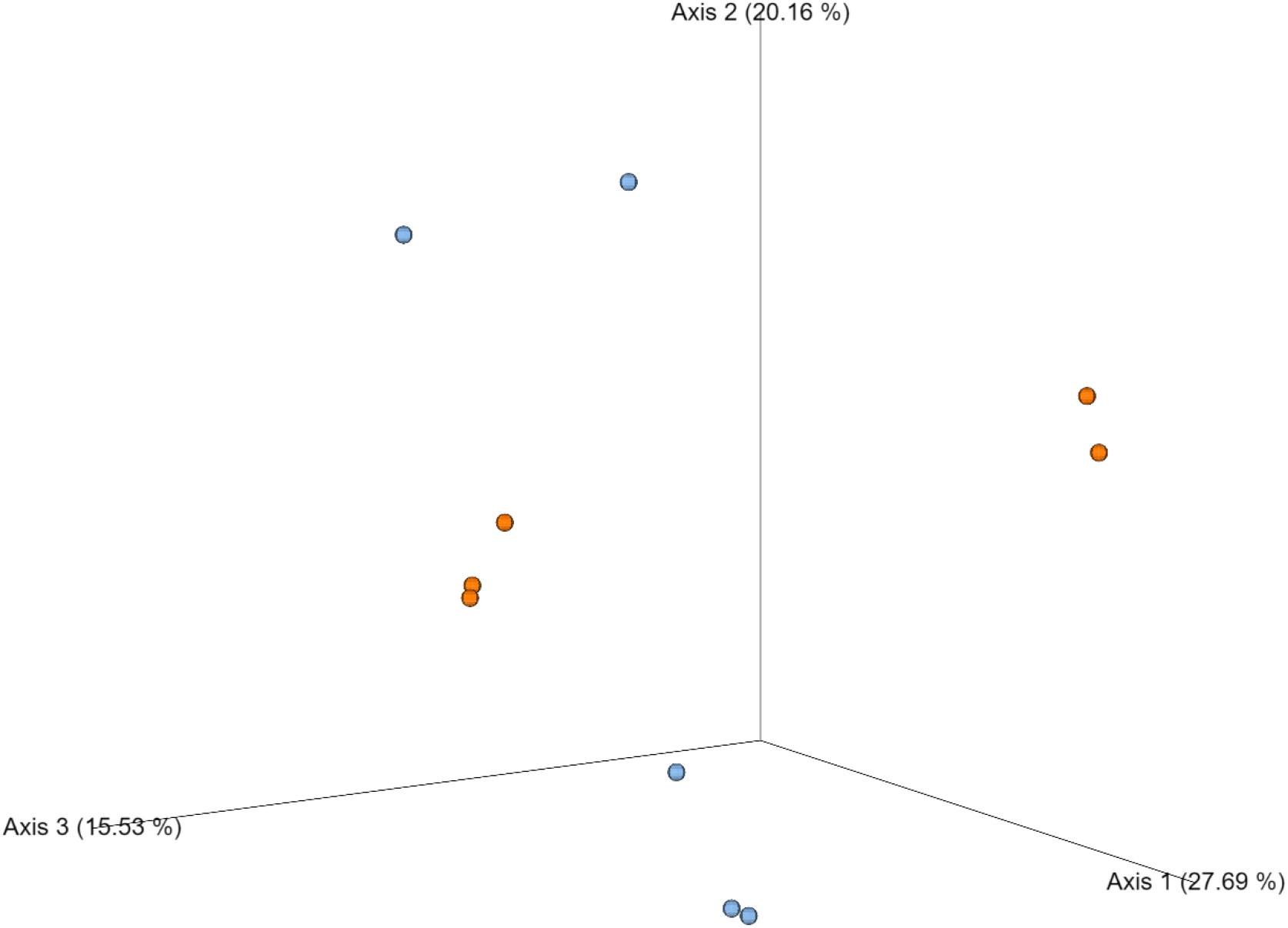
PCoA plot from Bray Curtis dissimilarity matrix. Percentage of the total variance represented by each axis is provided within parenthesis next to the label of the axis. Samples belonging to the experimental and control groups are indicated by orange and blue dots, respectively.

### 4. Correlation between questionary data and nasal microbiome

Concerning the alpha diversity, no significant differences in nasal microbiome composition were found in relation to sex, age, smoking habits and alcohol consumption. Nevertheless, females, non-smokers, age percentile 67-100% and individuals consuming alcohol *“once per week”* presented higher Shannon index and ASV abundance.

As far as the consumption frequencies of food related to different flavors is concerned, no significant differences in the nasal microbiome composition emerged when comparing individuals within and between groups, nor individuals taking part in different counts of tastings sessions over the last two weeks or consuming different types of wine.

The Spearman test evidenced a negative correlation between *“Smoking Habits”* and the presence of *Sphingomonas* (*χ^2^test; p*=0.007), *Xanthobacteraceae* (*χ^2^test; p*=0.025), *Radioerbium* (*χ^2^test; p*=0.013), *Pseudomonas* (*χ^2^test; p*<0.001) and *Micrococcus* (*χ^2^test; p*=0.035). Likewise, a negative correlation was found between *“Consumption frequency of bitter foods”* and *Corynebacterium* (*χ^2^test; p*= 0.005) and *Lawsonella* (*χ^2^test; p*=0.045) as well as between *“Consumption frequency of astringent food items”* and *Sphingomonas* (*χ^2^test; p*=0.048). Conversely, a positive correlation was found between *“Higher sensibility to sweet taste”* and the occurrence of both *Pelomonas* (*χ^2^test; p*=0.007) and *Caulobacteraceae* (*χ^2^test; p*=0.044), but the same enquired variable was negatively correlated with *Brachybacterium* (*χ^2^test; p*=0.013) and *Campylobacter* (*χ^2^test; p*=0.013). *“Higher sensibility to sour taste”* was positively correlated with *Pelomonas* (*χ^2^test; p*=0.007) and negatively correlated with *Haemophilus* (*χ^2^test; p*=0.046). *“Frequency of tasting liqueur wines”* was positively correlated with *Haemophilus* (*χ^2^test; p*=0.005) and *Veillonella* (*χ^2^test; p*=0.014) but negatively correlated with *Dolosigranulum* (*χ^2^test; p*=0.026). Negative and positive correlations were found between *“Frequency of tasting calm wines”* with *Corynebacterium* (*χ^2^test; p*=0.041) and *Cutibacterium* (*χ^2^test; p*=0.041), respectively. “*Frequency of tasting sparkling wines*” was negatively correlated with *Xanthobacteraceae* (*χ^2^test; p*=0.044).

Negative correlations were also found between *“Higher consumption of chili pepper”* and *Sphingomonas* (*χ^2^test; p*=0,028), *Brevibacterium* (*χ^2^test; p* <0.001), *Variovorax* (*χ^2^test; p*0.023), *Neisseria* (*χ^2^test; p*=0.025), *Haemophilus* (*χ^2^test; p*=0.025), *Prevotella* (*χ^2^test; p*=0.019), *Veillonella* (*χ^2^test; p*=0.025), *Enhydrobacter* (*χ^2^test; p*=0.013) and *Micrococcus* (*χ^2^test; p*=0.035). Individuals with a higher *“Higher consumption of lemon”* showed a negative and positive correlation with *Lactobacillus* (*χ^2^test; p*=0.045) and *Lawsonella* (*χ^2^test; p*=0.036), respectively. Two more positive correlations were found between *“Higher consumption of vinegar”* and *Porphyromonas* (*χ^2^test; p*=0.011), and *“Higher consumption of sugar”* and *Corynebacterium* (*χ^2^test; p*=0.008) (Figure 6).

**Figure 6.**
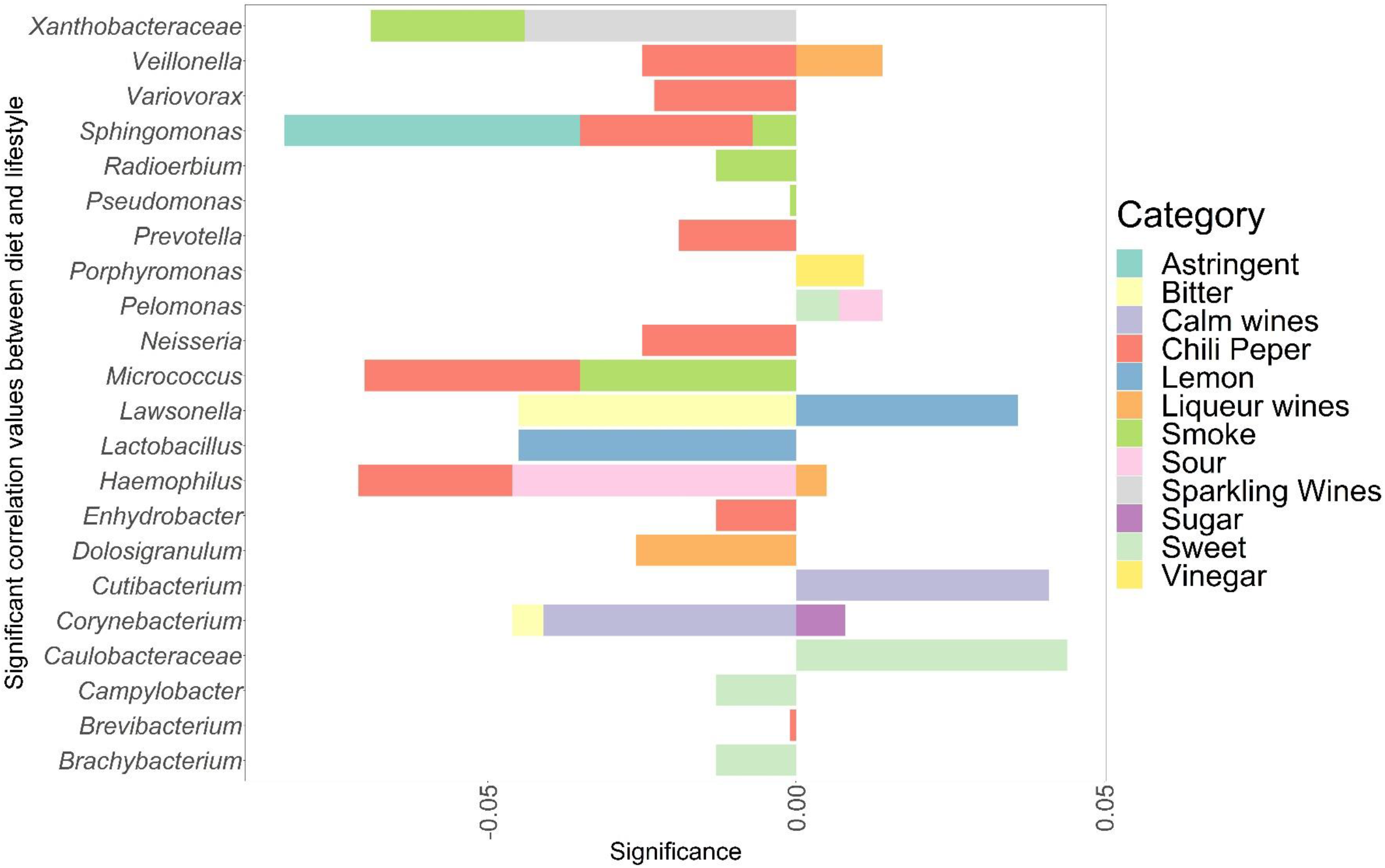
Horizontal barplot representing the correlation between each identified taxon of the nasal microbiome and the enquired variables related with diet and lifestyle. Correlation values [-1,1] are shown on the x-axis, where each bar represents the level of correlation between the taxon and the variables, right side bars indicate significant positive correlation and left side bars indicate significant negative correlation. The respective variable of each color bar are indicated in the legend: Higher sensibility to bitter food (Bitter); Higher sensibility to sweet food (Sweet); Higher sensibility to sour food (Sour); Higher sensibility to astringent items (Astringent); Smoking habits (Smoke); Tasting frequency of calm wines (Calm wines); Tasting frequency of sparkling wines (Sparkling wines); Tasting frequency of liqueur wines (Liqueur wines); Higher consumption of chili pepper (Chili pepper); Higher consumption of lemon (Lemon); Higher consumption of sugar (Sugar); Higher consumption of vinegar (Vinegar). The taxonomic names of the taxa are given on the y-axis.

### 5. *In-Silico* functional composition

The predictions of the metagenomic functional composition are shown in Figure 7. A total of 45 pathways were significantly enriched (Welch’s t-test; *p*<0.05). Of these, 33 and 12 were significantly higher and lower, respectively, in the experimental group than in the control group (Whelch’s t-test; *p*<0.05). TCA cycle related KEGG pathways (“TCA cycle IV”, “TCA cycle VII” “TCA cycle VIII” and “incomplete reductive TCA cycle”) were significantly higher in the experimental group. This process is one of the principal sources for bacteria to generate energy (aerobic process).

**Figure 7.**
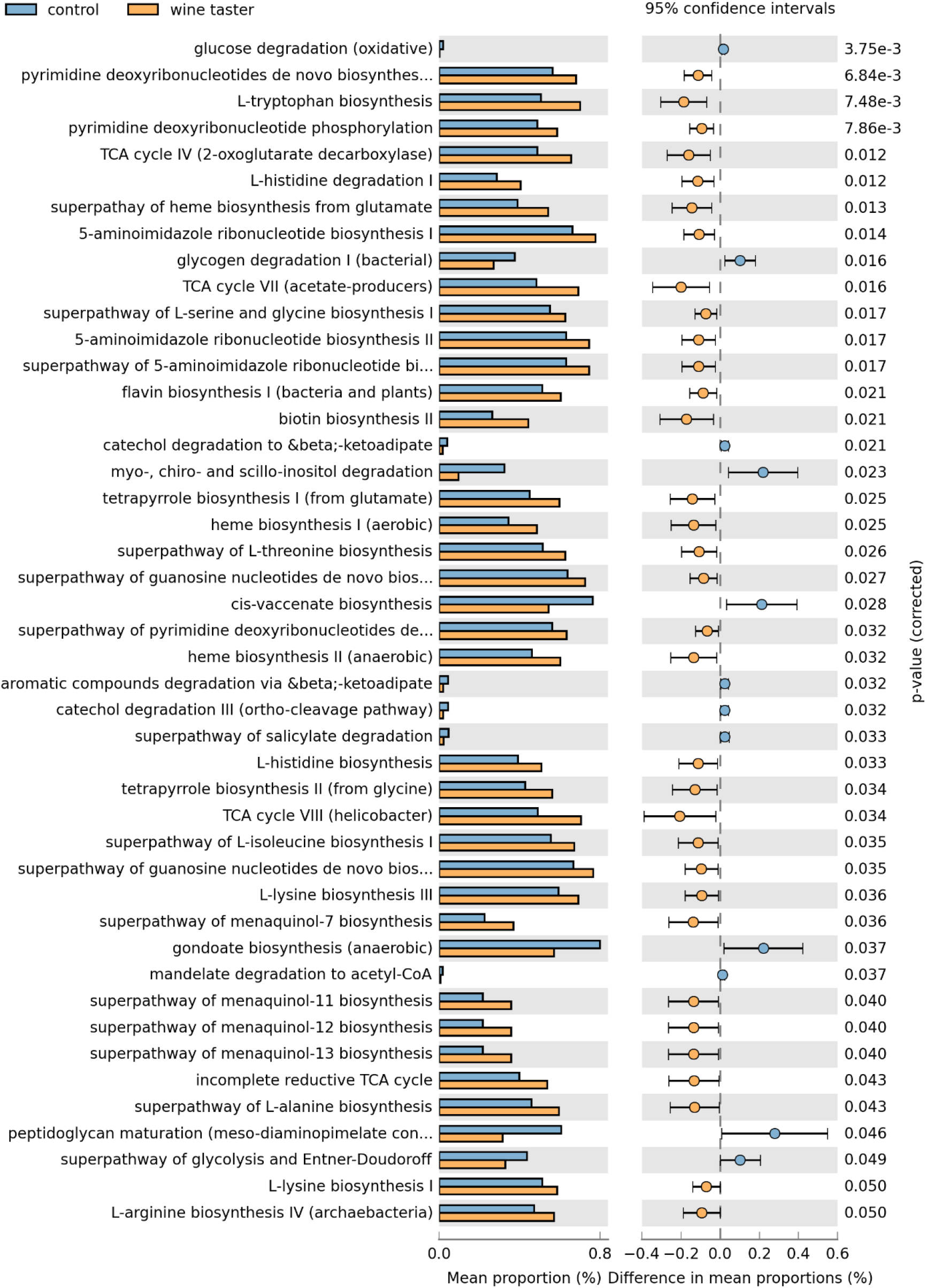
Predicted pathway abundances between the experimental and control group in the nasal microbiome. The KEGG database functional categories are indicated next to the displayed histograms (left panel: mean ± SD) and q-value determinations (right panel: 95% confidence intervals). Orange and blue colors denote individual cases of experimental and control groups.

## Discussion

This study aimed to characterize the nasal microbiota in wine tasters from Portugal using *16S rRNA* metabarcoding. Variables such as diet, oral hygiene, and lifestyle habits were also considered as they may influence this nasal microbiome.

Remarkably, there were no previous works on the nasal microbiome of the Portuguese population, nor specific studies in professional wine tasters. However, the taxa identified in our research, such as phyla Firmicutes and Actinobacteria as well as genera *Dolosigranulum* and *Corynebacterium,* had been previously found in similar works (Charlson et al., 2010; Lemon et al., 2010; Pfeiffer et al., 2022), which points to them as “normal” residents of the nasal microbiome.

Regarding the alpha diversity, the experimental group showed a lower SDI and ASV abundance when compared to the control group. However, the difference was not statistically significant. The genera *Brachybacterium, Lactobacillus, Granulicatella, Micrococcus* and family Neisseraceae were not found in the experimental group. Also, smokers showed a tendency for a decrease in microbiome diversity as opposed to previous results evidencing either a higher diversity in this category (Charlson et al. (2010) or no statistically significant differences with non-smokers Yu et al. (2017). Nevertheless, the finding of this latter study and the present one might be biased by the limited sample size. The negative correlation between smoking habits and the occurrence of five genera (*Sphingomonas*, *Xanthobacteraceae*, *Radioerbium*, *Pseudomonas*, and *Micrococcus*) deserves special interest and calls for the need of *ad hoc* studies based on larger samples sizes in order to address the consequences in the health and wellbeing. Another interesting preliminary finding in need of validation is the negative correlation between higher consumption of chili pepper and the presence of nine genera (*Sphingomonas, Brevibacterium, Variovorax, Neisseria, Haemophilus, Prevotella, Veillonella, Enhydrobacter*, and *Micrococcus*). It is worth mentioning, however, that a recent study evaluating the influence of a pepper-containing diet on the *Drosophila* gut microbiome revealed that variation in microbiome composition depends on animal genotype and pepper type (Garcia-Lozano et al., 2020). Studies with ample sample size and in humans need to validate the hypothesis that a pepper-containing diet could cause a shift in the nasal microbiome composition. These alterations could occur following the interaction with volatile chemicals inhaled through the nostrils. These compounds can also enter retronasally from the mouth when swallowing, contributing to the olfactory perception of food flavor (Dalton, 2004).

Concerning beta diversity, there was no significant difference between groups, indicating a substantial homogeneity of the nasal microbiome composition among individuals. The oral microbiome investigation performed on the same individuals (Duarte-Coimbra et al., 2023), showed higher microbiome diversity in the control group. Additionally, in that study, a larger number of taxa (497) was found (expected results since the oral microbiome is the second largest microbiome habitat in the human body after the gut microbiome, and that study presented a larger sample size). The nostril samples held a higher percentage of unassigned reads, a result totally expected since the nasal microbiomes have been far less studied than, for example, the oral ones.

The PCoA revealed one cluster composed by three individuals belonging to the experimental group and another by three individuals from the control group. Interestingly, the latter was also observed as one of the four aggregates of the hierarchical cluster, where samples tend to huddle with those of the same group, as originated by the abundance of *Dolosigranulum*, a genus including potential pathogens (Roitman et al., 2022).

Regarding bacterial functional prediction, we found that pathways involved in the Krebs cycle have a higher mean proportion among individuals of the experimental group, thus further evidencing a possible difference in the nasal microbiome composition of wine tasters. Higher exposure to alcohol molecules leads to many bacteria oxidizing ethanol into acetate by the pathway “TCA cycle VII – acetate-producers”. Subsequently, acetate enters the Krebs cycle as Acetyl-CoA simultaneously with coenzyme A and continues generating energy for bacteria (Wilson & Matschinsky, 2020).

The seemingly similar composition in the experimental and control group calls for caution due to the small sample size but represents a basis to build upon. When comparing the number of identified taxa and overall diversity, the group of wine tasters shows lower bacteria and a less diverse microbiome, thus possibly indicating a correlation between alcohol tasting and alterations in nasal microbiome composition.

## Conclusion

The present work was the first to analyze the nasal microbiomes in wine tasters and the first one to characterize this microbiome communities in the Portuguese population. Differences in the nasal microbiome diversity between wine tasters and non-wine tasters did occur, even though they were not statistically significant. Furthermore, there was a higher presence of Krebs cycle pathways in wine tasters, evidencing the importance of the nasal bacterial community in alcohol oxidation. These interesting preliminary results need validation in a larger sample size.

## Author Contributions

ABP, LPP and SDC designed the study; SDC executed the wet lab work under the supervision of GF; SDC and LPP performed the genomic data analysis; SDC drafted the MS and all authors contributed to the final version of the MS.

## List of Abbreviations

ASV: Amplicon Sequence Variant
PCR: Polymerase Chain Reaction
DNA: Deoxyribonucleic Acid
16S: rRNA 16S Ribossomal RNA
OTU: Operational Taxonomic Unit
QIIME: Quantitative Insights Into Microbial Ecology
RNA: Ribonucleic Acid
TCA: cycle VII Tricarboxylic acid cycle VII

## Acknowledgments

This work was supported by the European Union’s Horizon 2020 Research and Innovation Programme (Grant Agreement Number 857251) through the BIOPOLIS program. Lucía Pérez-Pardal and Giovanni Forcina are supported by national funds from Fundação para a Ciência e a Tecnologia and PTDC/BAA-AGR/28866/2017, respectively. The authors are thankful to Maria Magalhães (Centre for Molecular Analysis, CIBIO/InBIO) for her highly qualified laboratory work and assistance with library preparation.

